# Disruption of anterior temporal lobe reduces distortions in memory from category knowledge

**DOI:** 10.1101/2022.04.19.488831

**Authors:** Alexa Tompary, Alice Xia, H. Branch Coslett, Sharon L. Thompson-Schill

## Abstract

Memory retrieval does not provide a perfect recapitulation of past events, but instead an imperfect reconstruction of event-specific details and general knowledge. However, it remains unclear whether this reconstruction relies on mixtures of signals from different memory systems, including one supporting general knowledge. Here, we investigate whether the anterior temporal lobe (ATL) distorts new memories due to prior category knowledge. In this experiment (N=36), participants encoded and retrieved image-location associations. Most images’ locations were clustered according to their category, but some were in random locations. With this protocol, we previously demonstrated that randomly located images were retrieved closer to their category cluster relative to their encoded locations, suggesting an influence of category knowledge. We combined this procedure with transcranial magnetic stimulation (TMS) delivered to the left ATL before retrieval. We separately examined event-specific details (error) and category knowledge (bias) to identify distinct signals attributable to different memory systems. We found that TMS to ATL attenuated bias in location memory, but only for atypical category members. The magnitude of error was not impacted, suggesting that a memory’s fidelity can be decoupled from its distortion by category knowledge. This raises the intriguing possibility that retrieval is jointly supported by separable memory systems.

## INTRODUCTION

Our access to and use of semantic knowledge relies on the integrity of the anterior temporal lobe (ATL; Warrington 1975; Hart and Gordon 1990; Hodges et al. 1992). This knowledge is critical for forming and retrieving new memories of events, as even new experiences usually involve objects, places, and people for which we already have rich prior knowledge. Despite this, research on memory for events, or episodic memories (Tulving 1972), rarely considers the role of the semantic memory system, and in particular, the complexity and hierarchical organization of conceptual information in the formation and retrieval of new memories. What role does the ATL play in memories for new events that map onto a well-learned concept?

In past work, we developed an experimental protocol that aims to tease apart the fidelity of a memory and its influence by general knowledge—in this case, prior category knowledge—when retrieving the same encoded event (Tompary and Thompson-Schill 2021). We found that category knowledge systematically distorted episodic memories and we interpreted these findings through the lens of a memory reconstruction framework. In the current experiment, we modified this procedure for use with transcranial magnetic stimulation (TMS) to query the involvement of the ATL in these newly formed episodic memories. We addressed two questions: (1) whether disruption of ATL would result in a reduction in memory distortions, as predicted by memory reconstruction models, and (2) how this disruption may differentially impact new memories depending on their category typicality. Below we provide background for these two questions and our respective predictions.

### Memory reconstruction from multiple memory systems

Episodic memory has been well-characterized as a reconstruction of disparate sources of information, relying both on incomplete representations of the original event and relevant prior knowledge (Bartlett 1932; Huttenlocher et al. 1991; Hemmer and Persaud 2014). This integration process provides a good explanation for findings of enhanced memory for events that are consistent with prior knowledge (Bransford and Johnson 1972; Alba and Hasher 1983). However, such a reconstruction process comes at a cost for events that are inconsistent with prior knowledge. For instance, category knowledge often drives false memory creation (Deese 1959; Brewer and Treyens 1981; Roediger and McDermott 1995), and can produce small but systematic distortions in true memories (Hemmer and Steyvers 2009a; Hemmer and Steyvers 2009b; Persaud and Hemmer 2016; Brady et al. 2018). Such distortions are thought to be the product of an adaptive integration between prior knowledge and idiosyncratic details of the encoded event. Critically, prior knowledge and event-specific details are commonly found to be supported by distinct neural systems, raising the intriguing possibility that the retrieval process for a given memory may be supported by a mixture of signals from each. In the current experiment, we aim to understand whether brain regions in different memory systems provide neural signals that jointly support the retrieval of a single memory.

Several neuroscientific theories suggest that multiple brain regions may carry information from the same encoded event. For instance, Complementary Learning Systems (CLS; McClelland et al. 1995) posits that the anatomy of the hippocampus enables it to assign distinct, non-overlapping representations to overlapping inputs, such that new inputs can be rapidly learned without causing interference between memories. In contrast, cortex assigns overlapping representations to similar inputs, supporting learning of commonalities across multiple events. Building on this work, Trace Transformation Theory (TTT) proposes that over the course of systems-level memory consolidation, the shift in neural representation from the hippocampus to the cortex is accompanied by a transformation in what is remembered. Specifically, vivid, richly contextual memories continue to rely on the hippocampus, while more generalized memories are supported by cortex (Winocur et al. 2010; Sekeres et al. 2018). A central tenet of this model is that the brain stores both traces for the same event, and the relative strength of each trace dictates which is reinstated and in turn, how much specific detail versus generalized information is retrieved. In the current experiment, we hypothesized that ATL would fill the role of the cortical region representing more generalized memory, carrying information about the categorical organization of encoded images. Specifically, disruption of ATL through TMS would attenuate memory distortions arising from prior category knowledge, relative to performance in a control condition. Further, we predicted that the overall fidelity or precision of each location memory would remain unchanged, as this would likely be supported by the hippocampus, which was not disturbed.

### Category typicality in the anterior temporal lobes

Although CLS and TTT are largely agnostic about the cortical region that represents generalized knowledge, we targeted the ATL since we were interested in using category membership as a particular form of generalized knowledge in our experimental protocol. Converging evidence across patient work, neuroimaging, and causal methods has shown that this region supports the recognition, classification and production of common concepts (Warrington 1975; Snowden et al. 1989; Pobric et al. 2007; Binney et al. 2010; for a review, see Patterson et al. 2007). Of relevance to our experiment, damage to this region results in misclassification of the category membership of both manmade objects and living things (Hodges et al. 1995; Rogers et al. 2006; Rogers and Patterson 2007). Finally, transient disruption of the ATL through TMS has also revealed impairments in picture naming, object matching, and other tasks involving semantic processing of objects (Pobric et al. 2010a; Ishibashi et al. 2011; Chiou et al. 2013; Bonnì et al. 2015; Chiou and Lambon Ralph 2016; Woollams et al. 2017). These well-studied properties of the ATL make it a suitable target for our experiment objectives.

The second aim of this experiment was to investigate whether disrupting ATL would have a differential impact on memory distortions that could be predicted from the organization of their semantic elements. To do this, we leveraged the variation in typicality of members of a category, where typical category members share the greatest number of features with other category members (Rosch et al. 1976). Because of this internal organization of categories, typical items are more quickly categorized (Rips et al. 1973; Murphy and Brownell 1985), their features are more easily generalized to new exemplars (Rips 1975; Osherson et al. 1990), and they are more likely to be both correctly recalled (Schmidt 1996) and falsely recalled when excluded from a encoding list that includes members of the same category (Smith et al. 2000). Finally, research from patients with ATL damage consistently reveals a graded organization of semantic knowledge, such that patients are more likely to have access to more general or typical features of objects relative to more specific ones (Warrington 1975; Hodges et al. 1995). Although this property of semantic knowledge is robust and well-studied, it is less clear how category typicality and its neural basis influences the reconstruction of episodic retrieval. Thus, we included category typicality as a condition of interest in our protocol.

Given the strong evidence for category typicality as an organizing dimension of semantic knowledge, how exactly might disruption of the ATL differentially affect episodic memories involving typical and atypical category members? Findings from patient and TMS data suggest that patterns of error might become more similar to each other. This would indicate a ‘flattening’ of the category that is driven by a loss of knowledge about distinctive features, which would disproportionally affect atypical category members. This is suggested from observations of errors like mis-naming atypical category members as more typical ones—e.g. ‘horse’ for ‘zebra’—before reverting to its superordinate category name— ‘animal’ for ‘zebra’ (Hodges et al. 1995). Similarly, patients make drawing errors like taking away the distinguishing features of atypical category members and incorrectly adding features belonging to more typical category members—for instance, drawing a rhino without its horn or a duck with four legs (Bozeat et al. 2003; Patterson and Erzinçlioğlu 2008). Finally, in an object recognition protocol, TMS to ATL primarily affected typical category members, decreasing accuracy and slowing response times such that responses to typical category members more closely resembled those of atypical category members (Chiou and Lambon Ralph 2016). In the context of our experiment, these findings would generate the prediction that under TMS to ATL, the pattern of memory distortions for typical and atypical category members would become less distinct than we previously reported. However, another possibility is that inhibiting ATL will ‘contract’ the boundaries of a category, making it more difficult to associate more atypical items with their category. This pattern can be observed from semantic fluency findings, where patients with more advanced cases of semantic dementia become less likely to bring to mind more atypical category members in response to a category cue (Hodges et al. 1995). In the context of the current experiment, this would give rise to even less distorted memory for atypical category members than observed in participants with stimulation to a control region, and thus a larger difference in the extent of distortion relative between typical and atypical category members. Given these two conflicting predictions, we considered the differential impact of ATL disruption on memory distortions by category typicality as an exploratory analysis.

### Overview of experiment

In the present experiment, we tested whether TMS to ATL would reduce distortions in memory due to category knowledge. Specifically, participants encoded and retrieved image-location associations on a two-dimensional (2D) grid. Each image’s location was chosen such that most members of the same category (e.g., birds) were located near each other, but some typical and atypical category members were in random locations (Figure 1B). This configuration allows participants to learn that images from a certain category tend to cluster in a particular area as they encoded the locations of specific images. We calculated two measures of interest: error, a directionless measure of the fidelity of each image’s location memory, and bias, the proportion of error in the direction of an image’s category cluster. Importantly, error and bias could vary independently, such that memory for an image could be biased toward or away from its category cluster at the same level of error (Figure 2A). When previously using this protocol (Tompary and Thompson-Schill 2021), we found that when an image’s encoded location was far from its cluster of category neighbors, participants placed it closer to its category cluster at test. We further demonstrated that the category typicality (Rosch et al. 1976) of an encoded image explained the extent of this distortion in location memory.

**Figure 1.**
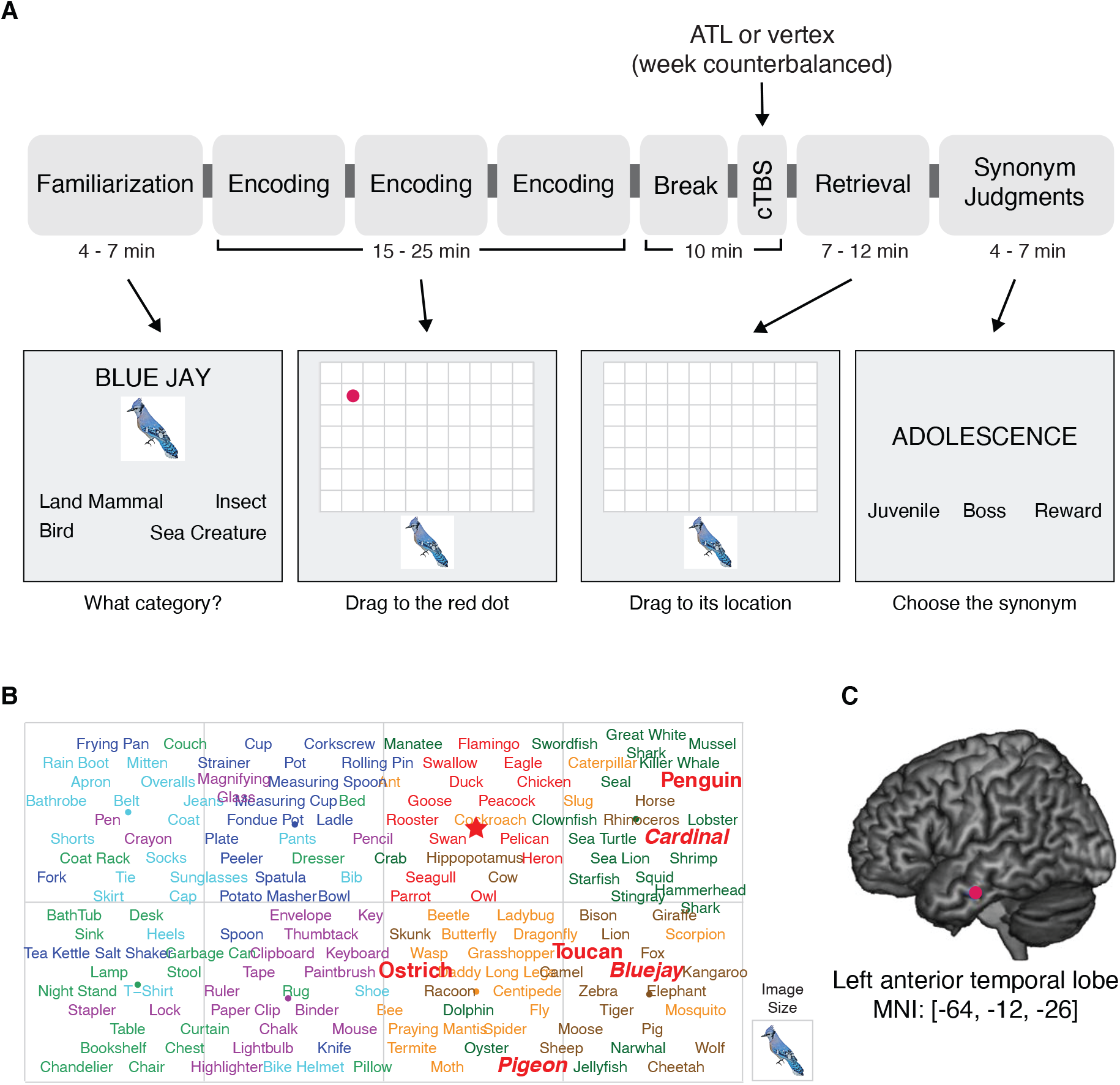
Experimental procedure. **(A)** Task order and design. Participants completed two sessions separated by one week, with stimulation to ATL or vertex counterbalanced. Each session included a familiarization task, three rounds of encoding in which all images were presented once, and then took a 10-minute break with cTBS administered in the last minutes of the break. Immediately after stimulation, participants completed a retrieval task and a synonym judgments task. **(B)** Example stimulus display for one participant. Each word corresponds to an image with that concept; font color indicates category membership (e.g. red words are birds). Image locations were divided into eight sections, or four per superordinate category. Spatially inconsistent words were randomly swapped into other sections on the same side of the screen (e.g. bolded red words indicate the locations of spatially inconsistent bird images). Half of the spatially inconsistent images were typical category members (italicized) and half were atypical category members (not italicized). **(C)** Site of stimulation targeting the left anterior temporal lobe. Red dot indicates MNI average coordinate across all participants.

**Figure 2.**
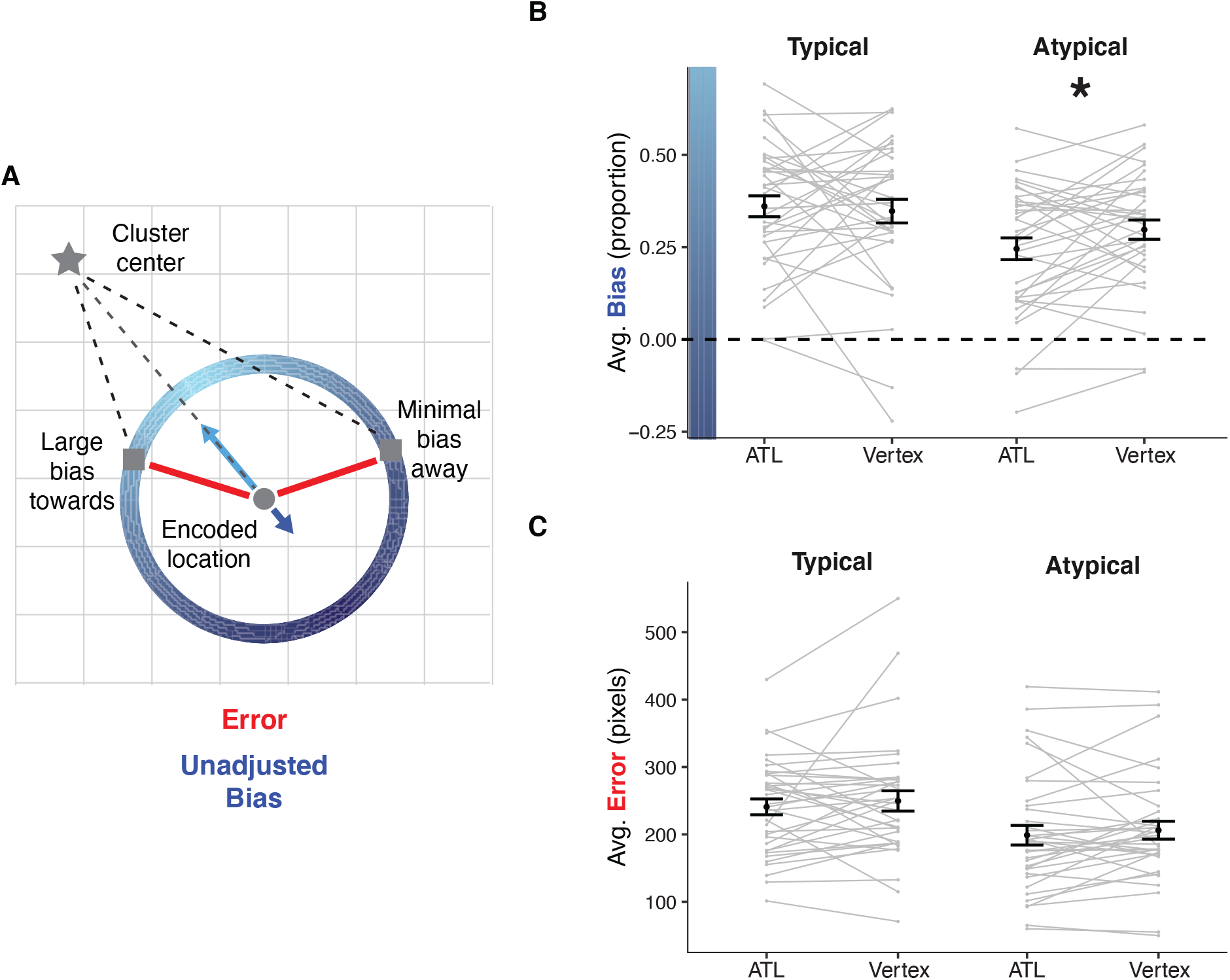
Analysis approach and results. **(A)** Examples of error and bias for two possible retrievals for the same image. Gray star indicates the center of an image’s category cluster. Gray circle indicates image’s encoded location. Gray squares indicate two possible retrieved locations. Red line indicates the magnitude of error, and blue errors indicate the extent of bias. Memory for an image could be biased toward or away from category neighbors at the same level of error, indicated by the blue circle. Shades of blue along the circle indicate different extents of bias for the same amount of error. **(B)** Average proportion of bias in location memory for typical and atypical category members. Dotted line indicates no bias towards or away from category clusters. * indicates *p* < .05. **(C)** Average error in location memory for typical and atypical category members. **(B-C)** Gray lines indicate participant averages. Black points indicate group average. Error bars indicate standard error of the mean across participants.

In the current experiment, participants completed this procedure in two separate sessions. In the experimental session, TMS to the left ATL was administered prior to retrieval, and in the control session, TMS was delivered to the vertex. We hypothesized that, under stimulation to vertex, we would replicate findings that memory for the locations of images is biased in the direction of their category’s general location. Specifically, memory for images of typical category members will be retrieved closer to other category members relative to images of atypical category members. Second, we hypothesized that the disruption of ATL via TMS will attenuate such biases in location memory. Third, we explored how the category typicality of the encoded items influenced the extent of ATL influence in their memory bias. Fourth, as most experiments that deliver TMS to ATL find that its disruption primarily impacts semantic processing, we included a synonym judgment task which has been used to demonstrate slowed semantic processing under TMS to ATL (Pobric et al. 2007, 2009). With this task, we aimed to replicate observations that synonym judgments are slower under stimulation to ATL relative to a control site.

## MATERIALS AND METHODS

### Participants

36 participants (20 female) ranging from 19 – 39 years of age (mean: 26 years) completed the experiment. We determined our pre-registered sample size based on a power analysis estimating the sample size needed to find our weakest predicted effect in the absence of stimulation (e.g. in our control condition). In a cohort of participants collected to validate our stimulus sets, we found that typical category members were retrieved closer to their category cluster relative to atypical category members, t(33) = 2.80, p = .009, replicating our prior findings (Tompary and Thompson-Schill 2021). A power analysis using this effect size (Cohen’s *d* = 0.48, alpha = 0.05, power = 0.8, two-sided, paired t-test) recommended a sample of 36 participants. We pre-registered a plan to exclude participants whose accuracy on the familiarization task was < 75%; no participant fell below that level of accuracy, thus no participants were excluded.

All participants were recruited from the University of Pennsylvania and greater Philadelphia area using online advertisements. Participants (1) were fluent English speakers, (2) reported no history of neurological impairments, (3) completed safety screening for TMS prior to the experiment. Participants were paid $100 upon completion of the experiment. The University of Pennsylvania IRB approved all consent procedures.

### Apparatus

TMS was delivered with a Magstim Super Rapid2 system with a figure-of-eight coil (70 mm). Positioning of the stimulation coil was guided using a frameless stereotaxic neuronavigation system (Brainsight 2, Rogue Research Inc.) paired with Polaris Vicra sensor camera and infrared reflecting markers that enabled registration between participants’ heads and their structural MRI. All tasks were coded in Javascript/HTML and were presented on a PC testing laptop.

### Materials

Stimuli for the memory tasks comprised 256 100×100 pixel color images on white backgrounds. Images were divided into two stimulus sets (Set 1 and Set 2). Each set contained two superordinate categories, each with four categories (Set 1 – *animals*: mammals, sea creatures, insects, and birds; *everyday objects*: kitchen utensils, office supplies, furniture, and clothes; Set 2 – *foods*: fruit, vegetables, grains, and seasonings; *objects requiring expertise*: sports equipment, construction tools, musical instruments, and vehicles). Note that ‘everyday objects’ and ‘objects requiring expertise’ were labels developed after stimulus development and do not perfectly capture distinctions between the two groups of stimuli. We had no a priori reason to separate objects based on this distinction but rather are using these labels as a shorthand way of labeling the different sets. Category typicality was determined with a list ranking procedure completed by a separate cohort of participants (Tompary and Thompson-Schill 2021).

In the memory tasks, all images were presented with an associated location on a white 600×1200 pixel rectangle with light gray gridlines spaced to form 50×50 pixel grids. To generate images’ locations for the memory tasks, the grid was divided into halves with one superordinate category on one side and the other superordinate category on the other side. On each side, all images were spaced uniformly apart, resulting in an even distribution of images across the entire grid. Each side’s locations were divided into four quadrants, and the four categories were randomly assigned to a quadrant (Figure 1B).

Then, the spatial locations of a subset of images were disrupted such that their locations were not consistent with category knowledge. To do this, images of the 3 most typical and 3 most atypical category members were swapped with the typical and atypical category members of other categories such that each quadrant included an equal number of typical and atypical category members from the other three categories. The remaining 10 images were randomly assigned to locations within their category’s quadrant. In total, 80 images were in locations that were consistent with their category membership (‘spatially consistent’), and 48 were in a random location (‘spatially inconsistent’). Of the 48 inconsistent images, 24 were typical and 24 were atypical category members. This procedure was conducted separately for the two stimulus sets.

Stimuli for the synonym judgment task were shared from prior investigations of the role of ATL in semantic processing (Pobric et al. 2007; Pobric et al. 2009). There were 144 target words, each paired with a synonym and two unrelated foils. The words are divided into two lists matched for frequency and imageability. The session in which each list was used was counterbalanced across participants.

### TMS procedure

Continuous theta burst stimulation (cTBS) was delivered in repeated trains of 200 bursts (3 50-Hz magnetic pulses per burst) with an inter-train interval of 200 ms (5 Hz), for a total of 600 pulses (40 sec). The stimulation was set at 80% of the resting motor threshold (RMT; Chiou et al. 2013; Chiou and Lambon Ralph 2016), separately for the two sessions. Motor threshold is defined as the minimum percentage of machine output required to produce motor evoked potentials (MEPs) of at least 50 μV on at least 5 of 10 consecutive trials at the same location. At this threshold, the average intensity of stimulation was 49% (SD = 8%) of the stimulator maximum output (range: 33% - 70%). Importantly, there was no difference in the intensity of stimulation when delivering TMS to ATL versus to vertex (*t*_(35)_ = 0.94, *p* = .35, *d* = 0.16). Six participants exhibited RMT that corresponded to a stimulation intensity that was too high for the machine to program; the stimulation was thus set to the maximum programmable intensity despite being lower than 80% of RMT. Excluding these subjects did not meaningfully change any results.

Using participants’ structural brain image, ATL was defined as the anterolateral region 10 mm posterior from the tip of the left temporal pole along the middle temporal gyrus (MNI: -53, 4, -32; Figure 1C) (Pobric et al. 2007; Lambon Ralph et al. 2009). The left ATL was chosen due to its prominent role in semantic processing in past TMS studies (e.g., Pobric et al. 2007; Pobric et al. 2009; Ishibashi et al. 2011; Chiou and Lambon Ralph 2016) although similar effects have also been found in the right hemisphere (e.g., Lambon Ralph et al. 2009; Pobric et al. 2009; Woollams et al. 2017). The control site vertex was defined as the midpoint between an individual’s nasion and inion, along the sagittal midline of the scalp (MNI: 0, •17, 65). MNI coordinates reflect approximate location, as all regions were defined separately for each participant based on anatomical landmarks.

### Experimental procedure

This study used within-subjects design with 1 factor (TMS site) and 2 levels (ATL and vertex). It comprised two sessions separated by 7 – 10 days. The majority were separated by 7 days unless there were constraints with the participants’ availability. The two sessions’ procedures were identical except for the site targeted by TMS and the stimulus sets used, both of which were counterbalanced across participants to create 4 counterbalancing groups. We used block randomization to ensure that an equal number of participants were allocated to each group (8 participants per group). The experimental procedure is identical to what we have reported previously (Tompary and Thompson-Schill 2021), except for an added familiarization task and a synonym judgments task. Each session is arranged in the following order: familiarization, encoding, 10-minute break with TMS stimulation, retrieval, and synonym judgments (Figure 1A). Following synonym judgments, participants completed two five-minute decision-making tasks: a risky decision task and temporal discounting task. Results from these tasks will be discussed in a separate manuscript.

#### Familiarization

This task served multiple purposes: (1) to introduce participants to the range of memoranda they would encode in the memory experiment; (2) to ensure equivalent categorization of the images across the stimulus sets, and (3) to exclude any non-compliant subjects. On each trial, participants viewed each image and four options. They were instructed to choose the option that best represents the image’s category. The options corresponded to the four categories that comprised the superordinate category of which the image was a member. For example, when viewing a cardinal, participants chose from bird, land mammal, sea creature, and insect, and when viewing a spatula, participants chose from kitchen utensil, office supply, furniture, and clothing. Participants used keyboard presses to indicate their choices. The mapping between options and keys were randomized for each participant. This task was untimed but participants were instructed to respond as quickly as possible while still being as accurate as possible.

#### Encoding

On each trial, participants viewed an image beneath the 600×1200 pixel grid and a red dot on the grid corresponding to that image’s location on the grid. They were instructed to drag each image onto the dot, click the mouse button or press the ‘enter’ key once the image was positioned over the dot, and try to remember each image’s location for a later memory test. Clicking the mouse automatically advanced the participant to the next trial. This task was the only task in the experiment that was not self-paced; if the participant did not move the item in under 6 seconds, the experiment automatically advanced to the next trial. All trials were presented a total of three times, in separate blocks, with the order of trials within blocks pseudo-randomized for each participant. The encoding instructions included two practice trials to familiarize participants with the task before beginning the first encoding block.

#### Retrieval

The retrieval task began immediately after stimulation. The timing and task were identical the encoding phase, but without a red dot marking the location of the image. Participants were instructed to drag the image to its location. After each retrieval trial, participants rated their memory for the image’s location as ‘Very confident’, ‘Somewhat confident’, ‘Guessed’, or ‘Forgot item’. Clicking on one option automatically advanced the participant to the next trial. The trial order was randomized for each participant.

#### Synonym judgments

On each trial, participants viewed a target word in the center of the screen and three words underneath: the synonym and the two unrelated foils. Participants were instructed to click on the synonym as quickly as possible while still being accurate. Trials were untimed and clicking on an option automatically advanced the participant to the next trial. The order of trials was randomized for each participant, and the order of the response options displayed on the screen were randomized on each trial.

### Measured variables

We used two dependent measures to assess memory for each image: error and bias due to category knowledge (Figure 2A). Both were developed previously (Tompary and Thompson-Schill 2021) and pre-registered for use in this experiment (https://osf.io/4j8vw/). Error was defined as the Euclidean distance between the encoded location and the retrieved location of an image, where greater values indicate less precision, and a value of 0 would correspond to perfect memory. Bias was defined as the proportion of error that is in the direction of an image’s category cluster. To do this, we first computed the unadjusted bias by subtracting the Euclidean difference between the encoded location and its cluster center from the Euclidean difference between the retrieved location and its cluster center. Then, we divided this unadjusted bias by the error for the image. Thus, a bias score of 0 indicates no bias, a score between 0 and 1 indicates that retrieval is biased towards the cluster center, and a score between 0 and -1 indicates that retrieval was biased away from the cluster center.

### Statistical models

All measures were entered into two-tailed paired t-tests and repeated measures ANOVAs. Wilcox rank sum tests were used in place of Student’s t-tests when data were not normally distributed, specifically for accuracy in the familiarization and synonym judgment tasks. We used an alpha of < .05 for determining significance in all statistical tests. Effect sizes are reported for all effects, including partial *η*^*2*^ for main effects or interactions of ANOVAs and Cohen’s *d* for effect sizes of within-subject comparisons. We calculated Cohen’s *d* as a within-subjects measure by incorporating the correlations across conditions (Lakens 2013), for easier comparison to our past work using the same experimental procedure (Tompary and Thompson-Schill 2021).

### Analyses – familiarization

Planned analyses for the familiarization included excluding participants who performed below 75% on this task, an extremely poor level of performance that would indicate non-compliance with the task. Accuracy was computed as the proportion of correct answers per participant. Across sessions and sites, performance on this task was consistently high (mean = 95.6%; SD = 3.7%), and no participants fell below the planned exclusion criterion. We also used this task to ensure equivalent categorization of the images across the two sessions, as the familiarization task took place before delivery of TMS. Because accuracy was near ceiling and thus not normally distributed, we computed a Wilcox ranked sum test over accuracy as a function of stimulation site. Accuracy on this task was not reliably different before delivery of TMS to ATL versus to vertex (*V* = 142, *p* = .16).

Although the main purpose of the familiarization task was to introduce participants to the range of memoranda they would encode in the memory experiment and to serve as an exclusion criterion, exploratory analysis of this data revealed effects of typicality that led us to modify our analysis of the memory experiment. Specifically, we found that relatively more typical category members were accurately categorized relative to atypical ones (*V* = 45.5, *p* < .001; Supplemental Figure 1). Errors in categorization of atypical category members often were for the second most likely category; for example, categorizing a penguin as a sea creature rather than a bird or categorizing a jet ski as sports equipment rather than a vehicle. Furthermore, log-transformed median response times were slower for atypical category members over typical ones (*t*_(35)_ = -11.42, *p* < .001, *d* = -1.9). Together, results from the familiarization phase indicate that participants were slower and less accurate when categorizing relatively more atypical category members compared to category members with high typicality, findings that fit with a long history of typicality effects in semantic processing (Murphy 2002; Patterson 2007). Because of this imbalance of categorization accuracy by typicality, when analyzing the retrieval task, we only included data from images that were correctly categorized.

### Analyses – memory

We pre-registered two analyses for the memory experiment: First, we planned to assess average error as a function of images’ consistency with prior knowledge, with a 2 (consistency: spatially consistent, spatially inconsistent) x 2 (site: ATL, vertex) ANOVA. We predicted that under stimulation to vertex, there will be more error for inconsistent images relative to consistent images, and that under stimulation to ATL, this difference in error would be diminished or eliminated.

Second, we planned to assess average bias amongst the inconsistently located images as a function of their category typicality, with a 2 (typicality: typical, atypical) x 2 (site: ATL, vertex) ANOVA. Our pre-registered hypothesis was that that under stimulation to vertex, typical category members would be more biased towards their category cluster relative to atypical category members, and under stimulation to ATL, this difference in bias would be diminished or eliminated. Because of the strong typicality effects present in the familiarization task, we chose to conduct exploratory analyses of bias by typicality by restricting analysis to items that were correctly categorized. We chose to do this because if participants were unable to correctly choose the category of an image, any influence of that image’s category cluster would be attenuated or nonexistent, diluting any possible influences of TMS on bias in the direction of its category cluster.

### Analyses – synonym judgments

We pre-registered one analysis for the synonym judgments. Here, we predicted slower response times under stimulation to ATL relative to vertex. We tested this by computing the median log-transformed response times of each participant separately for each site and entering these values into a two-tailed paired t-test. We computed this test including all trials regardless of accuracy, mirroring results published using the same stimuli (Pobric et al. 2007; Pobric et al. 2009). We additionally re-computed the analysis by excluding the first five trials of each session, trials with responses slower than 3 standard deviations from a participant’s median response time, and trials with responses faster than 100 ms. Although not pre-registered, we also include analyses of accuracy by TMS delivery for comparison to past work, calculating accuracy as the proportion of trials with correct responses and using a two-tailed paired Wilcox ranked sum test. We also expected that the disruption of ATL activity would influence multiple tasks requiring semantic processing. Therefore, we explored relationships between bias in location memory and performance on the synonym judgment task.

## RESULTS

### Bias by category typicality

A 2 (typicality: typical, atypical) x 2 (site: ATL, vertex) ANOVA including correctly categorized images revealed a main effect of typicality (*F*_(1, 35)_ = 21.65, *p* < .001, *η*^*2*^ = .40), replicating our previously published observation that typical category members are more biased towards their category cluster relative to atypical ones (Tompary and Thompson-Schill 2021). There was no reliable main effect of site (*F*_(1, 35)_ = 0.67, *p* = .42, *η*^*2*^ = .04). However, because this analysis revealed a trend for a typicality by site interaction (*F*_(1, 35)_ = 3.64, *p* = .07, *η*^*2*^ = .09), we conducted comparisons of bias by site separately for typical and atypical category members (Figure 2B). These paired t-tests revealed less bias after TMS to ATL relative to vertex, but only for atypical category members (*t*_(35)_ = -2.17, *p* = .04, *d* = -0.36) and not typical category members (*t*_(35)_ = 0.39, *p* = .70, *d* = 0.06). Surprisingly, TMS only impacted less typical category members, but the direction of this effect is in line with predictions from reconstruction model. Specifically, if TMS to ATL is disrupting its support of category knowledge, that would be result in less bias in memory towards the location of an image’s category cluster.

Note that the above analyses are constrained to category members whose images were correctly categorized in the familiarization phase (typical: mean = 97.5%, SD = 2.8%; atypical: mean = 92.8%, SD = 7.1%). We had pre-registered this analysis to use all trials regardless of categorization accuracy; the analogous 2 × 2 ANOVA including all trials revealed a main effect of typicality (*F*_(1, 35)_ = 24.32, *p* < .001, *η*^*2*^ = .41) and no main effect or interaction with site (both *F*’s < 1.84, both *p*’s > .18). We also conducted t-tests of the impact of TMS separately for typical and atypical category members, for closer comparison to the analysis of correctly categorized images. There was no reliable effect of site on bias (typical: *t*_(35)_ = 0.30, *p* = .76, *d* = 0.05; atypical: *t*_(35)_ = -1.51, *p* = .14, *d* = - 0.25). One potential reason TMS did not reliably impact bias here is that including incorrectly categorized images added noise to the dataset, diluting any subtle effects of TMS. This dilution would be extra strong for atypical category members, which were systematically less likely to be correctly categorized relative to typical category members.

### Error by category typicality

We conducted a similar 2 (typicality: typical, atypical) x 2 (site: ATL, vertex) ANOVA over the magnitude of error for all images that were correctly categorized. This revealed a main effect of typicality (*F*_(1, 35)_ = 52.81, *p* < .001, *η*^*2*^ = .66), again replicating our prior observations of greater error for typical over atypical category members. There was no reliable main effect of site (*F*_(1, 35)_ = 0.89, *p* = .35, *η*^*2*^ = .07) or interaction (*F*_(1, 35)_ = 0.02, p = .89, η^2^ = .001). Critically, TMS did not influence the magnitude of error for either typical or atypical category members (both *t*’s > -0.86, both *p*’s > .39, both *d’*s > -.14; Figure 2C). The same pattern of effects was found when following our pre-registered plan of including all trials.

### Bias by error for atypical category members

For atypical category members, TMS to ATL attenuated bias in memory due to category knowledge but not the overall magnitude error. This striking dissociation raised questions about the relationship between these two measures – for example whether disruption of ATL differentially impacted bias in memory such that only the weakest memories were impacted by TMS. To answer this question, we conducted a 2 (site: ATL, Vertex) x 4 (error: terciles 1 – 3) ANOVA across all atypical category members that were correctly categorized, where the error condition was computed by averaging bias for three equally-sized groups of images per participants, based on the magnitude of error for those images.

This revealed a main effect of site (*F*_(1, 35)_ = 4.92, *p* = .03, *η*^*2*^ = .03), echoing the impact of site on bias in atypical category members reported above. This ANOVA also revealed a main effect of error (*F*_(1, 35)_ = 5.87, *p* = .02, *η*^*2*^ = .09), such that regardless of TMS, images with a larger magnitude of error also exhibited larger biases towards their cluster center. This is consistent with the memory reconstruction framework: a memory whose item-specific representation is ‘noisier’ relies more on other, general knowledge, which in our case, would result in increased bias that comes from knowledge of the location of the image’s category cluster. Finally, this ANOVA revealed no reliable error-by-site interaction (*F*_(1, 35)_ = 0.03, *p* = .87, *η*^*2*^ = 0). In other words, the impact of TMS on bias did not change as a function of error. Instead, TMS to ATL resulted in less bias relative to TMS to vertex regardless of how accurately images were retrieved.

### Bias by superordinate category for atypical category members

In each session, participants learned the locations of two superordinate categories, one on each side of the grid. The use of four superordinate categories (two per session) enabled us to increase power while minimizing interference across trials and sessions, but it additionally presented an opportunity to identify whether the impact of TMS on bias was driven by a particular category. To this end, we computed an unpaired t-test comparing the effect of TMS on bias in atypical category members separately for the four superordinate categories (Supplemental Figure 2). Surprisingly, we found a reliable effect of TMS on bias only for animals (*t*_(24.6)_ = -2.40, *p* = .02, *d* = -0.82) and not for the other three superordinate categories (all *t*’s < 0<u>.35</u>, all *p*’s > .73, all *d*’s < .12). In other words, TMS to ATL only attenuated bias for images belonging to animal categories, not food or object categories. This effect does not seem to be driven by general differences in bias, since after TMS to vertex, bias for animals was not reliably different than bias for the other three superordinate categories (all *t*’s < 1.63, all *p*’s > .11, all *d*’s < .56).

### Error by consistency

The second class of analyses that we preregistered involved the relationship between spatial consistency and error, as we have previously found that images encoded far from their category clusters are less accurately remembered relative to images encoded within the cluster (Tompary and Thompson-Schill 2021). Through a reconstruction framework, the more accurate memory for spatially consistent images was interpreted as due to the additional ‘help’ of the category cluster in retrieving an image’s location. Here, we predicted that the difference in error by spatial consistency would be reduced after TMS to ATL, if this region indeed supports the category knowledge required to form knowledge of category clusters.

A 2 (spatial consistency: consistent, inconsistent) x 2 (site: ATL, vertex) ANOVA with error as the dependent variable, including all images that were correctly categorized. This model revealed a main effect of consistency (*F*_(1, 35)_ = 98.11, *p* < .001, *η*^*2*^ = .82), with more error in memory for spatially inconsistent images over spatially consistent ones regardless of TMS. This suggests that there is a strong influence of category knowledge on episodic memory, in that memory for spatially consistent images can draw on both details of the encoded event and category information that aligns with the event. This is a direct replication of previous findings (Tompary and Thompson-Schill 2021), the first to be administered in a lab setting rather than online, and extended to a new set of stimuli. There was no reliable main effect of site (*F*_(1, 35)_ = 0.16, *p* = .70, *η*^*2*^ = .02) or interaction between spatial consistency and site (*F*_(1, 35)_ = 2.72, *p* = .11, *η*^*2*^ = .07). The same pattern of effects was found when including all images regardless of categorization accuracy.

### Synonym judgments

Accuracy was high (mean = 94.3%, SD = 4.1%) and TMS delivery did not reliably influence the proportion of correct responses (*V* = 259, *p* = .84). Log-transformed median response times to synonym judgments did not reliably differ as a function of TMS, either when including all responses (*t*_(35)_ = -0.33, *p* = .75, *d* = -0.05) or when excluding outlier responses (*t*_(35)_ = -0.51, *p* = .61, *d* = -0.09). Finally, we explored whether response times in this task correlated with bias in memory, to test whether both tasks are supported by ATL, perhaps through related neural computations. Across individuals, the difference in median response times by TMS did not vary with the difference in extent of bias in atypical category members by TMS (*r*_(34)_ = -0.18, *p* = .28).

## DISCUSSION

In the current experiment, we delivered TMS to the left anterior temporal lobe (ATL) before retrieval of episodic memories to test the prediction that the ATL supports distortions in memory due to category knowledge without altering overall fidelity of the memories. Indeed, disruption of the left anterior temporal lobe (ATL) affected new memories that were encoded in an environment where category knowledge could aid new learning. Using a spatial location protocol, we first replicated prior results showing that memory of locations for images from the same semantic category was biased by their category membership. Specifically, locations of randomly placed images were retrieved closer to their category cluster relative to their encoded locations, and this bias in memory was weaker for atypical category members over typical ones. We found that TMS to ATL attenuated these biases in memory, but only for atypical category members and not for typical ones. Critically, TMS did not impact the magnitude of error in memory. Taken together, this is the first evidence of causal involvement of the ATL in biasing episodic memories through activation of prior category knowledge. Below, we situate these results within Trace Transformation Theory, offer some ideas for why the impacts of TMS were limited to atypical category members, and discuss some important caveats and avenues for future work.

Trace transformation theory (TTT) provides a compelling explanation for how learners can integrate relevant information from an event into networks of prior knowledge while also preserving episodic memory for its idiosyncratic details. This theory leverages anatomical distinctions between the hippocampus and cortex, as modeled by Complementary Learning Systems (McClelland et al. 1995), and posits that the brain stores both a hippocampal and a cortical trace to record the same event. The extent of reinstatement of each trace at retrieval may thus govern the amount of specific detail versus generalized information retrieved (Winocur et al. 2010; Sekeres et al. 2018). This raises the intriguing possibility that a memory’s retrieval is supported by *both* hippocampal and cortical signals, and variation in the strength of these signals has consequences for how the memory is expressed – a possibility we tested in the current experiment. We found that for atypical category members, disruption of the ATL attenuated bias in location memories. In other words, there was a reduction in participants’ tendency to retrieve image locations closer to other category members relative to where they were initially encoded (Figure 2B). At the same time, the magnitude of error in memory did not change (Figure 2C). This suggests that disruption of the ATL reduces the strength of neural traces that represent more generalized components of a memory but does not impact neural traces that represent its unique details. This uncoupling of error and bias further suggests that the retrieval of a single memory comprises different elements supported by discrete brain regions. In summary, our findings provide the first causal evidence that disrupting ATL function can reduce category bias but not episodic error in memory, suggesting that the retrieval of an encoded event comprises multiple elements which may map onto discrete memory systems.

One promising avenue for future work would be to test whether disruption of the hippocampus reduces error in location memories without impacting bias, thus preserving the influence of category knowledge while reducing the fidelity of the idiosyncratic details of each event. This would demonstrate a double dissociation of the contributions of hippocampus and cortex during memory retrieval, and thus bolster the claims of TTT. Already, there is some evidence that indirect stimulation to the hippocampus via a functionally connected cortical site can impact episodic retrieval (Wang et al. 2014; Hermiller et al. 2019; Hebscher and Voss 2020). Of particular relevance to this experiment, cTBS delivered to the hippocampus via the angular gyrus enhances the precision of location memories in a similar protocol (Nilakantan et al. 2017; Tambini et al. 2018), which raises questions of whether cTBS results in inhibitory or excitatory effects depending on the anatomy of the target site. Critically, to our knowledge, there have been no corresponding tests of more generalized memory that would demonstrate a selective impairment to memory for the episodic details of an event. There is some promising indication from autobiographical memory studies that disruption of episodic memory network regions reduces the number of internal details recalled in a memory, but either increase or do not impact the number of external details, which are thought to comprise more generalized and semantic information (Thakral et al. 2017; Bonnici et al. 2018). It is clear that more causal approaches are needed to fully test the hypotheses generated by TTT in humans.

Most consolidation theories either implicate the medial prefrontal cortex as a site of generalized knowledge or are agnostic to the exact cortical region that supports generalized memory traces (McClelland et al. 1995; Nadel et al. 2000; Winocur et al. 2010). We chose to target ATL due to its fundamental role in semantic processing. The logic behind this decision is that different cortical regions may store different types of generalized memory traces depending on their content or the salient features along which the category is organized. Because the generalized knowledge in the current experiment comprised spatial clusters that were linked together through their category membership, we predicted ATL might support this form of generalized memory. Interestingly, the ATL is particularly important for tasks requiring taxonomic category knowledge (Jefferies and Lambon Ralph 2006; Schwartz et al. 2009; Lewis et al. 2015). In other words, ATL is more likely to support categories that are organized by their attributes, (e.g. wings, fur) rather than by their function or relations (e.g. occupying the same contexts). This may explain why we observed the largest decrement in bias for the animal images, whose semantic organization is based more on attributes, relative to the object images, whose semantic organization is based more on shared relations or functions (Supplemental Figure 2). Disrupting a region like angular gyrus (AG), which is thought to represent concepts based on their shared functions (Binder et al. 2009; Boylan et al. 2017), may be more likely to result in reduced biases for those stimuli in the current experiment, a testable hypothesis for future work. Note however that because AG is functionally connected to the hippocampus (a connection that is crucial for the above-mentioned studies that deliver TMS to the hippocampus), it may be difficult tease apart its unique contribution. One way to do this may be to target its more anterior aspect, which is less functionally connected to the hippocampus relative to its more posterior aspect (Uddin et al. 2010).

Surprisingly, TMS to ATL reduced bias in memory for atypical category members but did not impact typical category members. Since atypical category members are generally less biased than typical ones, both in past behavioral work (Tompary and Thompson-Schill 2021) and when collapsing across stimulation site in the current experiment (i.e. the observed main effect of typicality on bias), the further attenuation of bias for atypical category members is in essence magnifying the difference in bias relative to typical category members. Why might TMS only impact memory for atypical category members, and what does it mean that this impact creates a larger distinction in memory by typicality? Our results suggest that, rather than a ‘flattening’ of a category such that the influence of a category on the retrieval of typical and atypical members becomes more equivalent, disrupting ATL ‘contracts’ the boundary of a category such that atypical category members are even less associated with their category than in a healthy brain. This is consistent with evidence that semantic dementia patients are less likely to produce examples of atypical category members when prompted with a cue (Hodges et al. 1995) and broadly consistent of the progression of semantic dementia in which patients first lose access to subordinate category members and atypical features while preserving superordinate knowledge (Warrington 1975). Since patients show more reliable memory for the general properties of objects than for their more specific features (Warrington 1975; Hodges et al. 1995; Done and Gale 1997) and often apply familiar or typical labels to semantically related objects (Hodges et al. 1995), it may be that the atypical information about categories is the type of information most likely to be more prone to impacts of TMS because it is relatively more fragile. It is also important to note that the effects of TMS in this experiment are equivalent to a limited, partial lesion of the ATL – both because the range of stimulation did not cover its full extent in the left hemisphere, and no disruption at all occurred in the right hemisphere. It is possible that with a larger extent of disruption, a pattern of effects more consistent with a ‘flattening’ account would have emerged. Such effects would be more consistent with later stages of SD in which patients lose access to broader category information, making errors that cross superordinate categories altogether (e.g. classifying an animal as an object) rather than selectively losing access to atypical category members in earlier stages (Hodges et al. 1995).

Already, we have discussed the notion that the fact that TMS delivers a partial, milder disruption relative to patients with more severe ATL damage may explain our observed pattern of effects, namely the attenuation of bias only in atypical category members and not typical ones, as well as the lack of impact on the magnitude error by spatial consistency. However, the timing of TMS in our experiment may also provide an explanation for these muted effects. According to reconstruction models, the integration of signals reflection event-specific details and more generalized knowledge occurs at the time of retrieval. If this is the case, the largest disruptions in generalized knowledge would occur in our current experimental protocol when TMS was delivered immediately before retrieval. However, an alternative possibility is that these sources of information are combined at encoding, leading to a single memory trace that is already distorted in space due to the presence of category knowledge as participants encode each image’s location. There is existing evidence that in this protocol, the utility of category knowledge during learning affects memory for the images, with better exemplar memory for atypical category members over typical ones – an effect that is eliminated when images are not clustered by category and is not easily explained by an account of reconstruction at retrieval (Tompary and Thompson-Schill 2021). Furthermore, drawing attention to category information can magnify differences in dimensions that explain category membership during a perception task that likely do not rely on retrieval computations (Goldstone 1994; Goldstone 1995; Livingston et al. 1998; Levin and Beale 2000). Delivering TMS before encoding may reduce participants’ access to category knowledge at the time of initial learning, leading to even less bias in location memory and perhaps also weakening the difference in memory accuracy for images located in their category cluster relative to images in random locations. Indeed, neuroimaging studies have revealed influences of prior knowledge both when encoding (van Kesteren, Fernández, et al. 2010; Tse et al. 2011; Bein et al. 2014) and retrieving (van Kesteren, Rijpkema, et al. 2010) new memories, suggesting that prior knowledge and event-specific details may be integrated at multiple points throughout the memory cycle.

TMS delivered to the anterior temporal lobe did not impact participants’ reactions times in a synonym judgement task, counter to several reports of slowed reaction times after disruption of ATL using the same word lists (Pobric et al. 2007; Pobric et al. 2009; Lambon Ralph et al. 2009). What might account for this failure to replicate past effects? First, the task may have been conducted too late to be impacted by the stimulation. Because retrieval was untimed and was always performed before the synonym judgment task, participants began the synonym judgment task approximately 5 – 14 minutes after TMS delivery, depending on the speed of their responses during retrieval. Although we chose to use a continuous theta-burst sequence due to its ability to suppress cortical excitability for up to 50 minutes, most of these estimates of the duration of TMS effects have been conducted in the motor cortex (e.g., Huang et al. 2005; Haeckert et al. 2021), impacting a system with anatomical differences could affect the temporal dynamics of TMS differently from that of the anterior temporal lobe. Furthermore, while inhibitory effects from cTBS are often observed for longer durations, the largest effects are observed within 5 minutes of stimulation (Chung et al. 2016). Second, although we used the same word lists as Pobric and colleagues, these stimuli were developed for use with British participants and thus included vocabulary that may have been slightly less familiar to the participants in our study. Although accuracy was near ceiling in and we found no difference in response times by site of stimulation when we excluded outlier responses, even a small increase in variability in response times may occlude or diminish any potential impacts of TMS. Third, the bulk of experiments that observed differences in response times with this protocol delivered TMS at 1-Hz pulses for 10 minutes, rather than a cTBS sequence (Pobric et al. 2007; Lambon Ralph et al. 2009; Pobric et al. 2009). Any of these possibilities may explain our failure to replicate; more work is needed to test the boundary conditions on the influence of TMS on ATL-dependent semantic processing.

One class of caveats for this experiment involves the limitations about our targeted region. First, the anterior temporal lobes are bilateral, and unilateral damage to these regions is less severe relative to bilateral damage or do not cause any semantic impairment (Hermann et al. 1999). Disrupting only the left hemisphere while leaving the right hemisphere intact may have attenuated the magnitude of bias in memory we observed or changed the nature of the bias completely. For example, perhaps disrupting both hemispheres would have lessened bias in memory for typical category members in addition to the observed reduction in bias only for atypical ones. Finally, it is worth noting that past experiments using TMS to disrupt either right or left ATL found equivalent deficits in semantic processing (Lambon Ralph et al. 2009; Pobric et al. 2010b), although when the task involves speech production such as in picture naming tasks, lateralized effects emerge in line with what would be predicted by the hemispheric lateralization of language networks (Woollams et al. 2017). Regardless, no study to our knowledge has disrupted both at once, leaving unanswered the question of how bilateral disruption would impact bias in memory. A second caveat involves ambiguity about the depth of the reach of our stimulation delivery – specifically, whether our stimulation protocol additionally impacted the anterior aspect of the left hippocampus in addition to the ATL. This possibility seems unlikely as the reach of TMS diminishes precipitously as a function of the depth into cortex, with stimulation from figure-of-8 coils able to achieve a maximum depth of about 3.4 cm (Deng et al. 2013) which falls short of the depth of the anterior hippocampus relative to the surface of the skull.

We have put forth a causal demonstration that the left anterior temporal lobe biases the retrieval of episodic memories, which are formed in conjunction with knowledge of category information. We believe this work offers an opportunity to better understand how information supported by both episodic and semantic memory systems is integrated in the context of new learning. These findings provide new insight into cognitive models of memory reconstruction by connecting them to neuroscientific theories of multiple memory systems and by addressing nuances related to the complex and rich organization of our semantic knowledge.

## ACKNOWLEDGMENTS

The authors are grateful for the members of the Thompson-Schill lab for useful feedback on the manuscript. We also thank Abby Clements for assistance with data collection, as well as Olufunsho K. Faseyitan and other LCNS lab members for technical assistance with the TMS equipment. Finally, many thanks to Dr. Gorana Pobric for sharing word lists for the synonym judgment task. Correspondence concerning this article should be addressed to Alexa Tompary, Department of Psychology, University of Pennsylvania, 3710 Hamilton Walk, Philadelphia, PA 19104, United States. Email: atompary@ sas.upenn.edu

## DATA AND CODE AVAILABILITY

Stimuli, raw data, and analysis code that support the findings of the TMS experiment will be made available in an OSF repository upon acceptance of the manuscript as a journal article. Raw data and analysis code dedicated to piloting and stimulus development will be made available upon request.

