## Supplemental Data for "Disruption of anterior temporal lobe reduces distortions in memory from category knowledge"

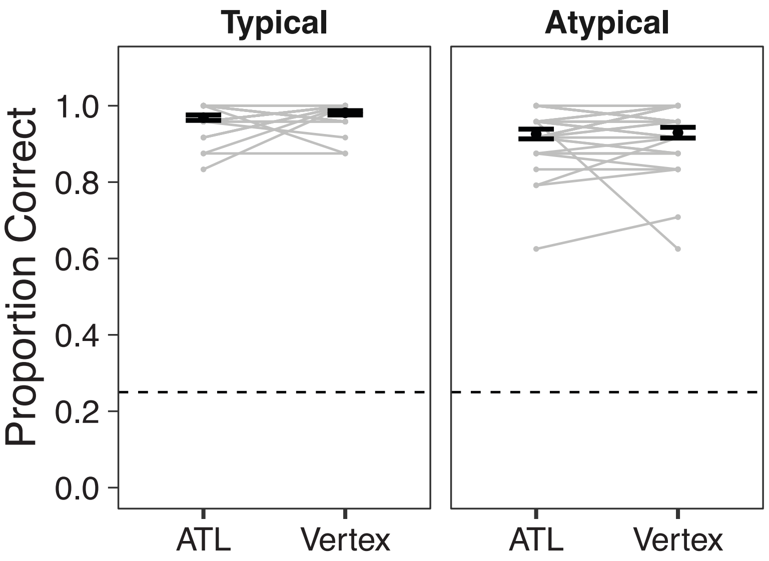


**Supplemental Figure 1.** Proportion correct responses in the familiarization task for typical and atypical category members. Gray lines indicate participant averages. Black points indicate group average. Error bars indicate standard error of the mean across participants. Dotted line indicates chance performance.

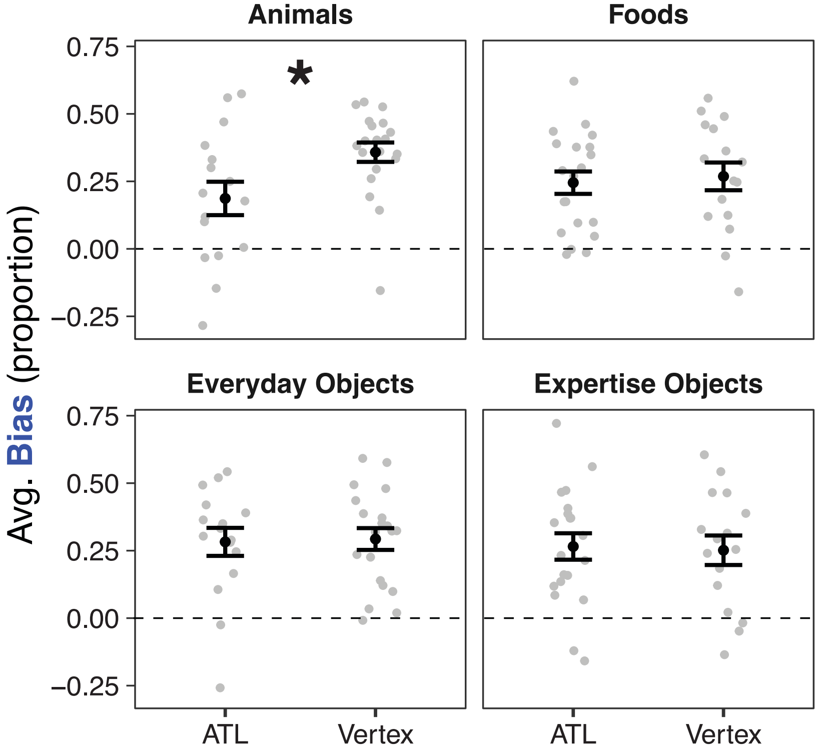


**Supplemental Figure 2.** Average proportion of bias in location memory for atypical category members, separately for the four superordinate categories. Note that ‘everyday objects’ and ‘objects requiring expertise’ are labels developed after stimulus development and do not perfectly capture distinctions between the two groups of stimuli. Gray dots indicate participant averages. Black points indicate group average. Error bars indicate standard error of the mean across participants. Note that all comparisons are between participants, as each participant interacted with separate superordinate categories across TMS sessions to reduce interference across them. Dotted line indicates no bias towards or away from category clusters. * indicates $p < .05$.
